# Overcoming rapaprotin resistance through inhibition of P-glycoprotein

**DOI:** 10.1101/2025.09.28.679086

**Authors:** Qian Zhang, Thomas Asbell, A. V. Subba Rao, Kalyan Kumar Pasunooti, Jian Zhang, Matthew G. Rees, Jennifer A. Roth, Jun O. Liu

**Affiliations:** Department of Pharmacology & Molecular Sciences, Johns Hopkins School of Medicine, Baltimore, MD 21205, USA; Broad Institute of MIT and Harvard, Cambridge 02142 MA, USA; Department of Oncology, Johns Hopkins University School of Medicine, Baltimore, MD 21205, USA; SJ Yan and HJ Mao Laboratory of Chemical Biology, Johns Hopkins University School of Medicine, Baltimore, MD 21205, USA

**Author notes:** These authors contributed equally (Qian Zhang, Thomas Asbell). Corresponding Author: Jun O. Liu Department of Pharmacology & Molecular Sciences Johns Hopkins University School of Medicine 725 N. Wolf Street, Hunterian 516 Baltimore, MD 21205.

**Keywords:** P-glycoprotein, Rapaprotin, Rapaprotin-L, DLD-1, substrate, drug resistance, 3D spheroid

## Abstract

The 26S proteasome is an essential regulator of protein homeostasis and a clinically validated therapeutic target in multiple myeloma (MM). Rapaprotin, a novel macrocycle identified from a rapamycin-inspired rapafucin library, disrupts 26S proteasome function by inducing disassembly of the 19S regulatory particle in the 26S proteasome, leading to apoptosis in MM cells. Its bioactivation requires prolyl endopeptidase (PREP)–mediated cleavage to generate Rapaprotin-L, a negatively charged, linear metabolite with potent proteasome-disassembly activity. Using the PRISM cancer cell line profiling platform, we identified high P-glycoprotein (P-gp/ABCB1) expression as a major determinant of Rapaprotin resistance in solid tumor cell lines. Efflux assays confirmed Rapaprotin-L, but not its parent Rapaprotin, as a high-affinity P-gp substrate. Co-treatment with the third-generation P-gp inhibitor tariquidar restored the intracellular accumulation of Rapaprotin-L, reinstating proteasome inhibition and consequent apoptosis of Rapaprotin-resistant colorectal cancer cell lines. Strong synergy between Rapaprotin and tariquidar was observed in a 3D spheroid model. These results establish P-gp as a key mediator of resistance to Rapaprotin and identify a rare example of a negatively charged Rapaprotin-L as a P-gp substrate. Together, these findings expand the potential therapeutic scope of Rapaprotin beyond hematologic malignancies to a broader range of solid tumors.

## Introduction

The 26S proteasome is a multi-subunit complex responsible for regulated protein degradation, a process critical for maintaining cellular homeostasis (Bard et al., 2018; Tanaka, 2009). It consists of a 20S catalytic core particle that mediates substrate proteolysis and a 19S regulatory particle that recognizes ubiquitin-tagged substrates, removes the ubiquitin chains, unfolds substrates, and translocates them into the 20S core for degradation (Bard et al., 2018; Tanaka, 2009). The proteasome is an attractive target for diseases with a high demand on the proteasome, such as multiple myeloma (MM) (Holkova and Grant, 2012; Hungria et al., 2019). This proteasome-dependent vulnerability of MM and mantle cell lymphoma has been successfully exploited through the development of FDA-approved 20S specific inhibitors, including bortezomib (BTZ), carfilzomib (CFZ), and ixazomib (IZB) (Holkova and Grant, 2012; Hungria et al., 2019). Despite their clinical efficacy, however, these 20S targeted proteasome inhibitors suffer from inevitable drug resistance and several adverse effects, particularly neurotoxicity (Thibaudeau and Smith, 2019; Yang et al., 2021). We recently identified Rapaprotin as a novel proteasome inhibitor from our rapamycin-inspired novel macrocycle rapafucin library (Fig. 1A). Rapaprotin operates via a novel mechanism--it induces the disassembly of the 19S regulatory particle from the 26S proteasome complex (Guo et al., 2019; Peng et al., 2025). This disruption leads to inhibition of ubiquitin-dependent degradation of substrates by the 26S proteasome, causing selective cytotoxicity in MM cells (Peng et al., 2025). Moreover, Rapaprotin requires activation by prolyl endopeptidase (PREP), which cleaves its inactive cyclic form to generate Rapaprotin-L; an active, linear, and charged form (Peng et al., 2025). Unlike conventional proteasome inhibitors that block catalytic activity, Rapaprotin exhibits selectivity for MM cell lines over primary endothelial cells and circumvents certain classic resistance mutations in PSMB5 and PSMB8 in bortezomib-resistant cells (Lu and Wang, 2013; Shi et al., 2020).

**Figure 1.**
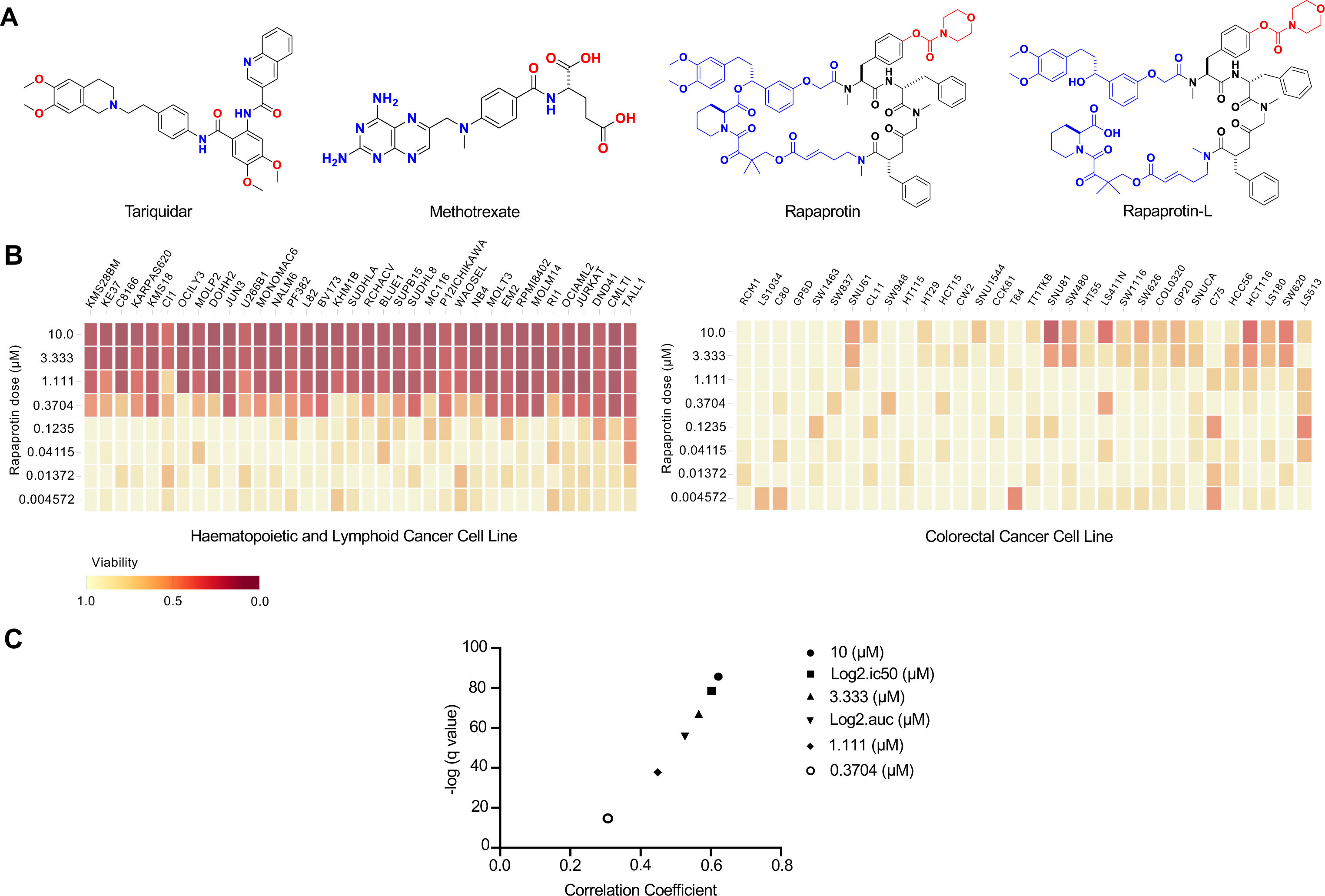
Positive correlation between ABCB1 (P-gp) expression and rresistence to Rapaprotin. **(A)** Structures of Tariquidar, Methotrexate, Rapaprotin, and Rapaprotin-L. **(B)** Heat map of cellular viability of Haematopoietic, Lymphoid, and Colorectal cancer cell lines treated with Rapaprotin. **(C) -**Log(q) values vs Pearson correlation of a high-throughput cell viability assay using over 900 genetically characterized human cancer cells suggests a positive correlation between ABCB1 (P-gp) expression and Rapaprotin treatment resistance. Coefficients and q values were calculated for each corresponding dose of Rapaprotin.

To further explore the potential use of Rapaprotin in a wide variety of cancer types, we performed the PRISM cancer cell viability profiling (Corsello et al., 2020). Analysis of drug-response data from a collection of over 900 cancer cell lines revealed a significant correlation between high P-glycoprotein (P-gp, MDR1, ABCB1) expression and resistance to Rapaprotin, suggesting that Rapaprotin and possibly its activated linear Rapaprotin-L may be substrates for P-gp.

P-gp is a 170-kDa ATP-dependent efflux transporter belonging to the ATP-binding cassette (ABC) family (Ahmed et al., 2022; Fu, 2013). It is highly expressed in barrier tissues such as the intestinal epithelium, liver canalicular membrane, kidney proximal tubules, and the blood–brain barrier, where it plays a protective role by transporting a wide range of xenobiotics and metabolic byproducts out of cells (Ahmed et al., 2022; Fu, 2013; Pilotto Heming et al., 2022). However, this same function becomes a major impediment in oncological drug discovery, as P-gp is frequently overexpressed in various cancer types and contributes to multidrug resistance (MDR) by actively exporting chemotherapeutic agents out of malignant cells (Pilotto Heming et al., 2022). High P-gp expression is associated with resistance to traditional chemotherapies, poor prognosis, and treatment failure in multiple cancers, including breast (Clarke et al., 2005), glioblastoma (Pilotto Heming et al., 2025), ovarian (Nanayakkara et al., 2018), and colon cancer (Beklen et al., 2020) amongst many others. This transporter reduces intracellular accumulation of structurally and mechanistically diverse anticancer agents such as taxanes (e.g., paclitaxel, docetaxel), vinca alkaloids (e.g., vincristine), anthracyclines (e.g., doxorubicin), and tyrosine kinase inhibitors (e.g., gefitinib, imatinib, and cediranib), thereby conferring broad-spectrum chemotherapy resistance (Pilotto Heming et al., 2022). Structural studies have shown that the P-gp binding pocket comprises a hydrophobic outer region and a deeper polar/charged region, which together accommodate a diverse array of substrates (Aller et al., 2009; Martinez et al., 2014).

Pharmacologic inhibition of P-gp has been explored as a strategy to reverse drug resistance and resensitize tumors to chemotherapies. First-generation inhibitors such as verapamil and cyclosporine A demonstrated modest success in early clinical studies but were hindered by low specificity, suboptimal affinity for P-gp, and significant off-target effects leading to unwanted toxicity (Amin, 2013; Belpomme et al., 2000; Ozols et al., 1987). Second-generation inhibitors, such as valspodar and dexverapamil, improved on these limitations but both failed due to either a lack of clinical efficacy or unwanted CYP450 interactions (Amin, 2013; Kolitz et al., 2010; Kovarik et al., 1998; Nguyen et al., 2021). More recently, third-generation inhibitors such as tariquidar (TQR) have shown higher specificity and greater potency, with reduced pharmacokinetic interactions. TQR has demonstrated promising results in Phase I and II trials as a safe and effective modulator of P-gp function when co-administered with chemotherapeutic agents (Amin, 2013; Fox et al., 2015; Pusztai et al., 2005).

Given the strong correlation between ABCB1 expression and resistance to Rapaprotin, we hypothesized that Rapaprotin, or its PREP-bioactivated Rapaprotin-L, is a potential substrate for P-gp. In this study, we tested this hypothesis using a number of cancer cell lines with varied levels of P-gp expression. Moreover, we investigated whether the P-gp inhibitor TQR can restore sensitivity to Rapaprotin in resistant cell lines, which may provide a strategy to broaden the anticancer activity to a wider variety of cancer cell types, especially those from solid tumors.

## Materials and Methods

### Reagents and antibodies

Reagents used for cell culture, including fetal bovine serum (FBS), penicillin/streptomycin solution, Trypsin-EDTA solution, Hank’s Balanced Salt Solution (HBSS), and GlutaMAX™-I, were obtained from Gibco (USA). Resazurin was purchased from Sigma-Aldrich, and the P-glycoprotein (P-gp) inhibitor tariquidar/XR9576(Cat. #S8028) was acquired from Selleckchem (USA). The following primary antibodies were used: anti-K48-linked polyubiquitin (rabbit polyclonal, Cell Signaling Technology [CST], Cat. #4289), anti-PARP (rabbit polyclonal, CST, Cat. #9542), anti-IκB-α (rabbit polyclonal, CST, Cat. #9242), anti-GAPDH (mouse monoclonal, Santa Cruz Biotechnology, Clone 0411, Cat. #sc-47724), and anti-P-glycoprotein (mouse monoclonal, Clone C219, Invitrogen, Cat. #MA1-26528).

### Cell Culture

DLD-1, A549, HeLa, MCF-7 and SW480 cells were cultured in Dulbecco’s modified Eagle’s medium (DMEM) supplemented with 10% FBS and 1% penicillin/streptomycin at 37°C with 5% CO_2_. NCI-H929 cells were cultured in RPMI-1640 medium and HCT116 cells were cultured in McCoy’s 5A medium with the addition of 10% FBS and 1% penicillin/streptomycin at 37°C with 5% CO_2_. All cell lines were purchased from ATCC.

### PRISM Profiling Assay

The PRISM cancer cell viability profiling was performed by Broad Institute (Cambridge, MA). Rapaprotin was treated at 8 doses in triplicate across 900 PRISM barcoded cancer cell lines for five days at the Broad Institute (Corsello et al., 2020). All cell lines were cultured in RPMI 1640 with 10% FBS. Adherent cell lines were seeded at 1250 cells per well and suspension cell lines were seeded at 2000 cells per well in pools between 20-25 cell lines per pool. Pools of cell lines were lysed with Qiagen TCL buffer to isolate the mRNA. mRNA was then reverse-transcribed into cDNA and then the sequence containing the unique PRISM barcode was amplified using PCR. Finally, Luminex beads that recognize the reverse complement of the specific barcode sequences in the cell set were hybridized to the PCR products. The magnetic beads attached to the PCR products were then detected using a Luminex FlexMap 3D scanner, which reports signal as a median fluorescent intensity (MFI). The data was then processed to calculate AUC, IC_50_, and create dose response curves. The details of the data processing steps are at: https://github.com/cmap/dockerized_mts

### Western Blotting

Western blotting (WB) was performed as described previously (Chen et al., 2022). Cells were washed and harvested in phosphate-buffered saline (PBS) prior to lysis in 1× RIPA buffer [0.1% SDS (w/v), 0.5% sodium deoxycholate (w/v), and 1% Nonidet P-40 (v/v) in PBS] supplemented with protease and phosphatase inhibitor cocktails. Lysates were centrifuged at 12,000 × g for 20 minutes at 4 °C to remove debris. Protein concentrations in the supernatants were determined using the bicinchoninic acid (BCA) assay and diluted in SDS loading buffer. Equal amounts of protein (10–20 µg) were separated by SDS–PAGE and transferred to polyvinylidene fluoride (PVDF) membranes (Millipore). Membranes were blocked, then incubated with primary antibodies followed by HRP-conjugated secondary antibodies. Protein bands were visualized using enhanced chemiluminescence (ECL) and imaged on a GeneSys Image Station. Densitometric analysis was performed using ImageJ. Primary antibodies were diluted in TBS containing 1% Triton X-100 and 5% BSA.

### Alamar Blue Cell Viability Assay

This assay was performed as previously described (Peng et al., 2025). Cells were seeded in 96-well plates (Corning) at the following densities in 190 μL media: NCI-H929, 2×10^4^ cells/well; DLD-1, A549, HeLa, MCF-7 and SW480, 750 cells/well. Subsequently, drugs were added at appropriate dilutions to achieve a final concentration of 0.1% DMSO. For adherent lines, drugs were added after an overnight incubation to allow cell attachment. After 72 hours of drug treatment, 10 µL of Alamar Blue reagent (Sigma) was added to each well, followed by incubation at 37°C for an additional 5 h. Fluorescence was then measured using a plate reader at 544 nm excitation / 590 nm emission. GraphPad Prism (v9.5.1) was used to analyze the data and determine IC_50_ values using a four-parameter logistic regression model. Synergy analysis was performed using SynergyFinder+.

### Measurement of Intracellular Concentrations of Rapaprotin and Rapaprotin-L

The intracellular accumulation of Rapaprotin and Rapaprotin-L was determined as previously reported (Peng et al., 2025). DLD-1 cells were seeded in a 6-well plate at a density of 5×10^5^ cells/well in 1 mL DMEM medium with 10% FBS, and allowed to adhere to plate overnight at 37 °C with 5% CO_2_. Rapaprotin and Tariquidar were added into the cell culture to a final concentration of 1 μM and 250 nM, respectively. Cells were incubated for designated time periods. At each treatment time point, cells were harvested, washed twice in PBS, and lysed in 50 μL of 1x RIPA buffer. Rapaprotin and Rapaprotin-L were extracted from cell lysate or 50 µL of cell culture media by adding 50 μL of saturated ZnSO_4_ and 150 μL of acetonitrile (ACN), vortexing for 30 seconds and centrifuging for 5 mins. The acetonitrile phase was separated and analyzed using HPLC-MS (Agilent 6120 Single Quad) with a Selective Ion Monitoring (SIM) mode. HPLC was run at a flow rate of 1.0 mL/min with a gradient of acetonitrile (with 0.1% formic acid) from 40-95% in water (with 0.1% formic acid). Mass intensity was calculated using the total peak area for m/z 1334.6 (Rapaprotin + H^+^) and m/z 1356.6 (Rapaprotin + Na^+^) for Rapaprotin and that for m/z 1352.6 (Rapaprotin + H2O + H+) and m/z 1374.6 (Rapaprotin + H2O + Na+) for Rapaprotin-L.

### Identification of P-gp Substrates

The assessment of Rapaprotin and Rapaprotin-L as potential P-gp substrates was performed by Pharmaron (Ningbo, China) using MDCKII-MDR1 cells in a bidirectional permeability format.

#### Preparation of MDCKII-MDR1 Cells and Assessment of Cell Monolayer Integrity

MDCKII-MDR1 cells were seeded on HTS Transwell Permeable Support inserts (0.4 μm pore polycarbonate membrane, 4.26 mm diameter, 0.14 cm^2^, 96-well culture plates) (Corning 3391) and allowed to expand to a final density of 5.45×10^5^ cells/cm^2^ in 4-8 days in an incubator at 37 °C with 5% CO_2_ and 95% relative humidity. Following replacement of medium from each Transwell insert and reservoir, the cell monolayer integrity was assessed by measuring transepithelial electrical resistance (TEER) across the monolayer using the Millicell Epithelial Volt-Ohm measuring system (Millipore). The plates were returned to the incubator once the measurement was completed. The TEER value was calculated according to the following equation:

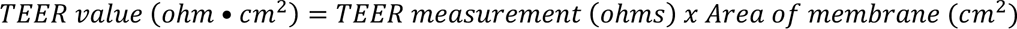

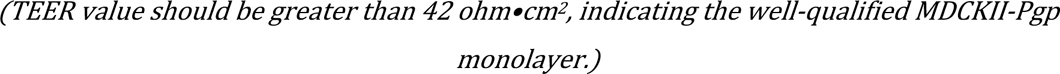

#### Compound Transport Assay

MDCKII-MDR1 cell monolayer was washed twice with pre-warmed HBSS (10 mM HEPES, pH 7.4). The plate was incubated at 37 °C for 30 minutes. To determine the rate of drug transport in the apical to basolateral direction, 125 µL of 1 µM test compound (diluted with HBSS (10 mM HEPES, pH 7.4)) was added to the apical compartment, followed by the immediate transfer of 50 µL samples to 200µL of acetonitrile containing IS (100 nM alprazolam, 200 nM caffeine, 200 nM labetalol and 100 nM tolbutamide) in a new 96-well plate as the initial donor sample (A→B). Samples were then vortexed at 1000 rpm for 10 minutes. Wells in the receiver plate (basolateral compartment) were filled with 235 µL of transport buffer (HBSS, with or without 10 µM PSC833). To evaluate transport in the basolateral-to-apical (B→A) direction, a separate 96-well plate was used for the initial donor sample (B→A), and the corresponding receiver wells were similarly prepared.

At the end of a 2-h incubation at 37 °C, 50 μL samples from both the donor sides (apical compartment for Ap→Bl flux, and basolateral compartment for Bl→Ap) and receiver sides (basolateral compartment for Ap→Bl flux, and apical compartment for Bl→Ap) were transferred to wells of a new 96-well plate. Four volumes of acetonitrile containing IS were then added. Samples were vortexed for 10 minutes and centrifuged at 3,220 × g for 40 minutes. An aliquot of 100 µL of the supernatant was mixed with an appropriate volume of ultrapure water prior to LC-MS/MS analysis. To assess Lucifer Yellow leakage (an indicator of monolayer integrity) after the 2-hour transport period, 100 µL of 100 µM Lucifer Yellow solution (prepared in HBSS (10 mM HEPES, pH 7.4)) was added to each Transwell insert (apical compartment). Wells in the receiver plate (basolateral compartment) were then filled with 300 µL of HBSS (10 mM HEPES, pH 7.4). After incubation at 37 °C for 30 minutes, 80 µL samples were collected from both apical and basolateral wells (using the basolateral access holes) and transferred to a new 96-well plate. Lucifer Yellow fluorescence was measured using a fluorescence plate reader (excitation at 480 nm and emission at 530 nm).

#### Data Analysis

Apparent permeability (P_app_) can be calculated for drug transport assays using the following equation:

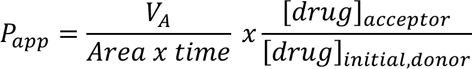

Where:

*P_app_* is apparent permeability (cm/s x 10^-6^);

*V_A_* is the volume (in mL) in the acceptor well;

*Area* is the surface area of the membrane (0.143 cm^2^ for Transwell-96 Well Permeable Supports);

*Time* is the total transport time (in seconds);

*[Drug]_acceptor_ and [Drug]_initial, donor_* are the concentrations of the drug in the acceptor and initial donor wells, respectively.

The efflux ratio was determined using the following equation:

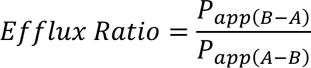

Where:

*P_app (B-A)_* is the apparent permeability coefficient in basolateral-to-apical direction;

*P_app (A-B)_* is the apparent permeability coefficient in apical-to-basolateral direction.

Mass balance (% recovery) was calculated using the following equation:

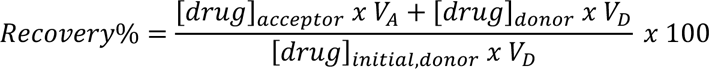

Where:

*V_A_* is the volume (in mL) in the acceptor well (0.235 mL for A→B flux, and 0.075 mL for B→A);

*V_D_* is the volume (in mL) in the donor well (0.075 mL for A→B flux, and 0.235 mL for B→A).

Lucifer Yellow (LY) leakage of the monolayer can be calculated using the following equation:

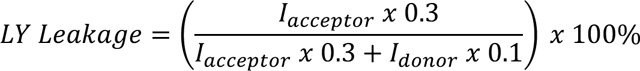

Where:

*I_acceptor_* is the fluorescence intensity in the acceptor well (0.3 mL);

*I_donor_* is the fluorescence intensity in the donor well (0.1 mL).

Lucifer Yellow leakage values should be less than 1.5% to indicate acceptable monolayer integrity. However, if the P_app_ values obtained from a specific well are qualitatively similar to those from replicate wells, based on scientific judgment, the monolayer may still be considered acceptable.

### 3D Spheroid Assay

Three-dimensional (3D) spheroid formation was performed using Corning Ultra-Low Attachment (ULA) 96-well microplates (ThermoFisher, Cat. No. 174925), as described previously (Sanchez et al., 2023). DLD-1 cells were seeded at a density of 1 × 10³ cells/well in ULA plates containing DMEM supplemented with 10% fetal bovine serum (FBS). Plates were then centrifuged at 400 × g for 5 minutes to promote cell aggregation. Cells were subsequently incubated undisturbed at 37 °C in a humidified atmosphere of 5% CO₂ for 72 hours to allow spheroid stabilization. Rptn and TQR were added at the indicated concentrations and incubated for an additional 3, 5, or 7 days, respectively. At each designated time point, 2 µM Calcein AM (Invitrogen) and 4 µM Propidium Iodide (PI) (ThermoFisher) were added to the spheroids for viability staining. After a 1-hour incubation at 37 °C, fluorescent images were acquired using a Nikon ECLIPSE TS100 microscope at the following excitation/emission settings: 494 nm/517 nm for Calcein AM and 493 nm/636 nm for PI.

### Statistical Analysis

Statistical analysis was done by the GraphPad Prism version 9.5.1 (GraphPad Software). All data are presented as mean ± standard error of the mean (SEM) and were analyzed by one-way ANOVA or two-way ANOVA, as appropriate. A p-value < 0.05 was considered statistically significant. Statistical significance is indicated as follows: *p < 0.05, **p < 0.01, and ***p < 0.001.

## Results

### P-Glycoprotein expression correlates with Rapaprotin resistance in cancer cell lines

Rapaprotin was identified from the rapafucin library using a proliferation assay of the multiple myeloma (MM) cell line NCI-H929, followed by structural optimization (Peng et al., 2025). Through inhibition of 26S proteasome function by dissociating 19S from 20S core particle, Rapaprotin induces apoptosis in MM cell lines. In contrast, it failed to induce apoptosis in HEK293T kidney epithelial cells or A549 lung cancer cells (Peng et al., 2025). To systematically assess the sensitivity of different types of cancer cells to Rapaprotin, we screened Rapaprotin in the PRISM collection of over 900 genetically annotated cancer cell lines at the Broad Institute (Corsello et al., 2020). As expected, MM cell lines are among the most sensitive subtypes (Fig. 1B). Importantly, a univariate gene-drug correlation analysis revealed a strong positive correlation between resistance to Rapaprotin and elevated expression of ABCB1, the gene encoding P-glycoprotein (P-gp) (Fig. 1C). This observation prompted us to investigate whether P-gp may play a role in the sensitivity of different cancer lines to Rapaprotin.

To validate this correlation, we analyzed baseline P-gp expression and Rapaprotin sensitivity in a panel of colorectal and non-colorectal cancer cell lines. Western blotting confirmed that DLD-1 and SW480 cells expressed high levels of P-gp protein, while A549, NCI-H929, HeLa, HCT116, and MCF-7 cells did not express detectable levels of P-gp (Fig. 2A-B). Correspondingly, DLD-1 and SW480 exhibited minimal sensitivity to Rapaprotin even at high concentrations, with <20% reduction in viability at 1 µM, whereas A549, NCI-H929, HeLa, HCT116, and MCF-7 cell lines were inhibited by rapaprotin in a dose-dependent manner with IC₅₀ values in the low nanomolar range (Fig. 2C-D). Together, these results strongly suggest that P-gp expression correlates with—and may contribute causally to—Rapaprotin resistance.

**Figure 2.**
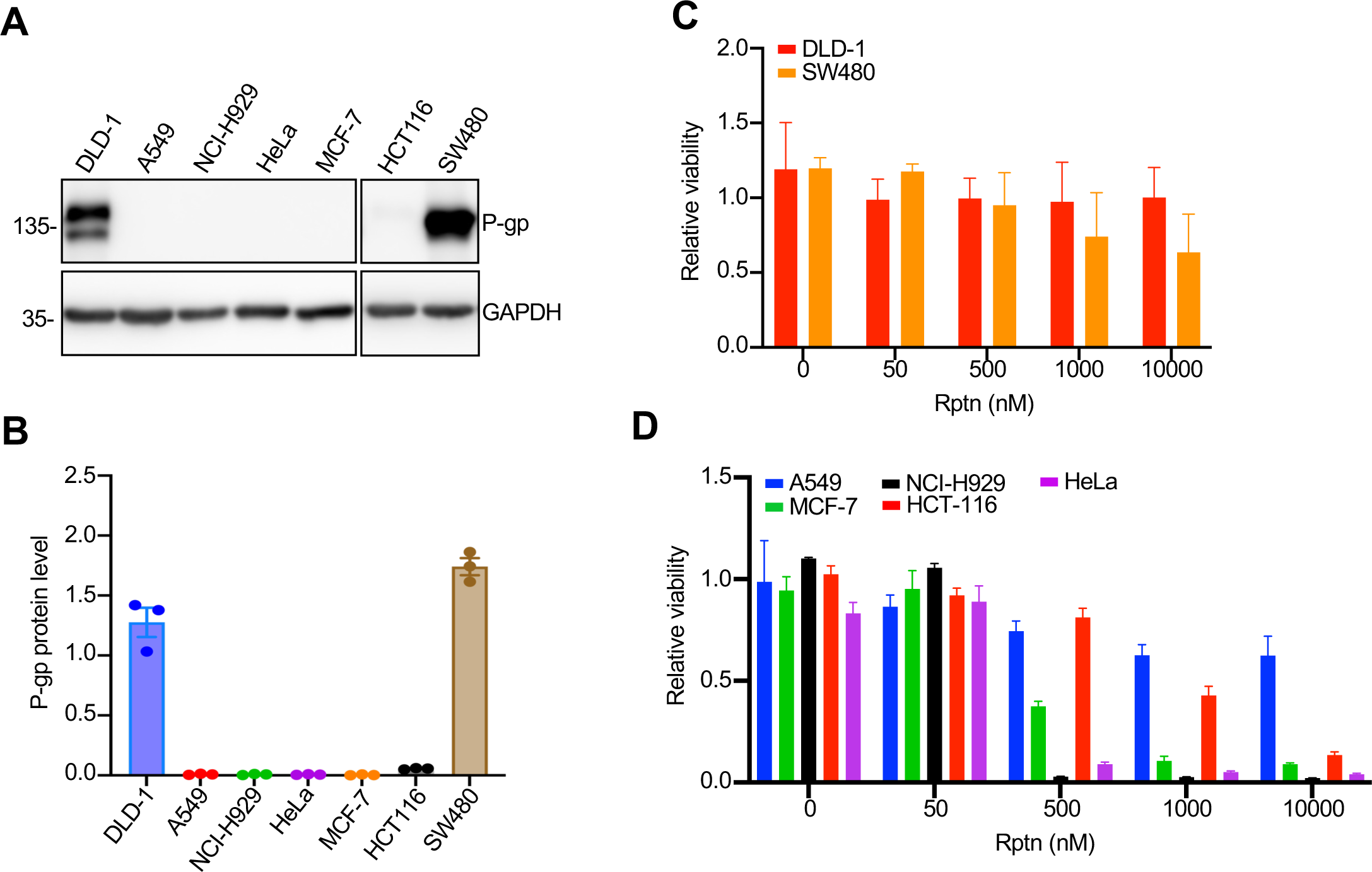
High P-gp expression correlates with Rptn resistance. (**A-B**) P-gp protein levels in DLD-1, A549, NCI-H929, HeLa, MCF-7, HCT116 and SW480 cell lines. (**A**) Cells were collected and lysed for Western blotting. (**B**) Quantitative analysis of protein levels in (**A**). Data were from three independent experiments and shown as mean ± SEM. (**C**) Relative cell viability of cell lines highly expressing P-gp. (**D**) Relative cell viability of cell lines sparsely expressing P-gp. Data were from three independent experiments and shown as mean ± SEM.

### Rapaprotin-L is a P-gp substrate and accumulates inside cells upon tariquidar cotreatment

We next asked whether P-gp directly effluxes Rapaprotin or its active metabolite, Rapaprotin-L, as a mechanism of resistance. As previously reported, Rapaprotin is a latent prodrug that requires intracellular processing by prolyl endopeptidase (PREP) to generate active Rapaprotin-L, which possesses potent proteasome-disrupting activity (Peng et al., 2025). Both rapamycin and FK506, which are structurally related to Rapaprotin, are known substrates of P-gp (Saeki et al., 1993; Yacyshyn et al., 1996; Yokogawa et al., 1999). Thus, it would not be surprising if Rapaprotin is also a P-gp substrate. Unlike Rapaprotin, Rapaprotin-L has a long linear structure containing a negatively charged carboxylic acid group. We have shown that Rapaprotin-L can be trapped in MM cells to reach high concentrations (Peng et al., 2025). Thus, we expected that Rapaprotin-L is less likely a substrate for P-gp.

To assess whether Rapaprotin and Rapaprotin-L are P-gp substrates, we conducted efflux assays in MDCKII-MDR1 cells stably overexpressing human P-gp. To our surprise, though Rapaprotin exhibited a moderate efflux ratio of 4.3, it remained unchanged in the presence of the P-gp inhibitor PSC833 (valspodar), suggesting a P-gp–independent efflux mechanism (Table 1A). In contrast, Rapaprotin-L displayed a strikingly elevated efflux ratio of 15.7, which was substantially reduced upon PSC833 co-treatment, indicating that Rapaprotin-L is a high-affinity substrate of P-gp (Table 1B). These findings provide mechanistic evidence that P-gp–mediated efflux of Rapaprotin-L, rather than the cyclic Rapaprotin, underlies Rapaprotin resistance.

**Table 1.** Permeability of Rapaprotin (A) and Rapaprotin-L (B) in MDCKII-MDR1 cell line

| Compound ID | PSC833 (μM) | Papp (A-B) ($10^{-6}$, cm/s) | Papp (B-A) ($10^{-6}$, cm/s) | Efflux Ratio | Recovery (%) A-B | Recovery (%) B-A |
| --- | --- | --- | --- | --- | --- | --- |
| Metoprolol | - | 28.53 | 29.07 | 1.02 | 117.0 | 101.66 |
| Digoxin | - | 0.46 | 9.07 | 20.06 | 104.92 | 94.83 |
|  | 10 | 1.43 | 1.90 | 1.33 | 101.87 | 91.61 |
| Rapaprotin | - | <0.03 | 0.14 | >4.31 | <107.27 | 11.04 |
|  | 10 | <0.03 | 0.13 | >4.29 | <113.84 | 66.52 |

(B)
| Compound ID | PSC833 (μM) | Papp (A-B) ($10^{-6}$, cm/s) | Papp (B-A) ($10^{-6}$, cm/s) | Efflux Ratio | Recovery (%) A-B | Recovery (%) B-A |
| --- | --- | --- | --- | --- | --- | --- |
| Metoprolol | - | 26.72 | 28.00 | 1.05 | 102.03 | 91.17 |
| Digoxin | - | 0.53 | 10.40 | 20.03 | 86.57 | 102.65 |
|  | 10 | 2.35 | 2.19 | 0.95 | 87.68 | 100.08 |
| Rapaprotin-L | - | <0.01 | 0.08 | >15.68 | <80.79 | 20.66 |
|  | 10 | 0.08 | 0.06 | 0.82 | 78.59 | 63.39 |

To validate this efflux in a cellular context, we determined intracellular concentrations of Rapaprotin and Rapaprotin-L in DLD-1 cells treated with either Rapaprotin alone or in combination with tariquidar. LC-MS analysis revealed a significant increase in intracellular Rapaprotin-L levels upon P-gp inhibition, while Rapaprotin levels remained unchanged, corroborating the observation made in the efflux assay (Fig. 3A-B). These results confirm that P-gp activity limits Rapaprotin efficacy by reducing the intracellular accumulation of its activated form that disassembles 26S proteasome, raising the possibility that cotreatment with tariquidar and other P-gp inhibitors can resensitize resistant cancer cells to Rapaprotin.

**Figure 3.**
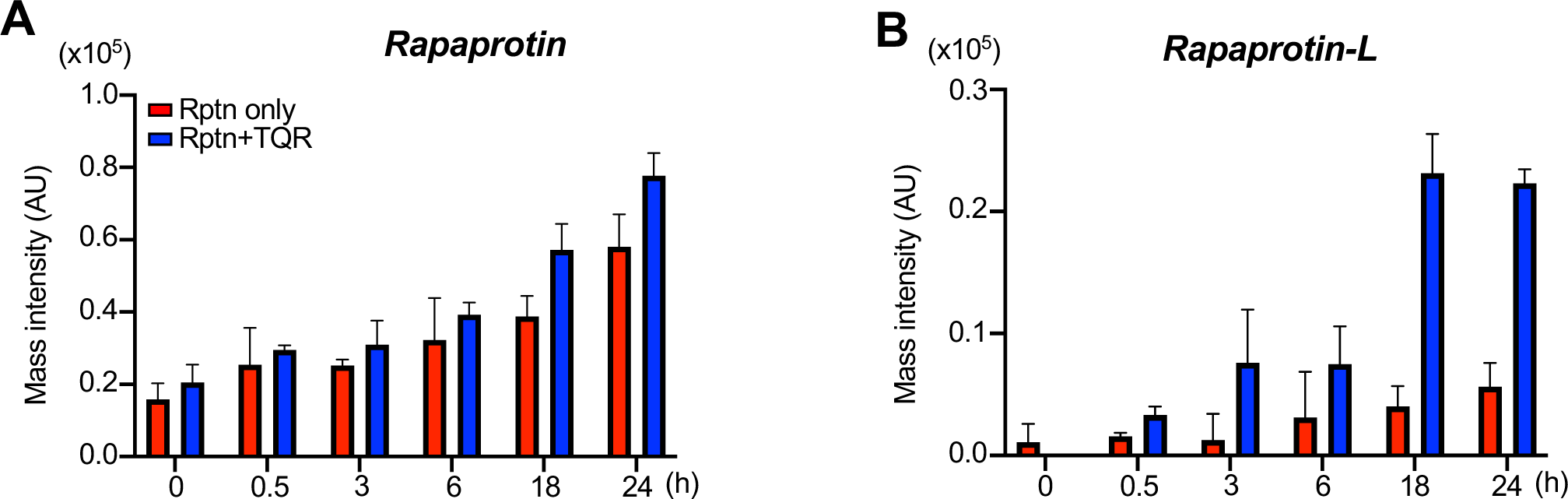
Rptn-L is a substrate of P-gp. (**A-B**) Mass intensity of intracellular Rapaprotin (**A**) and Rapaprotin-L (**B**) signals in DLD-1 cells treated with or without 150 nM Tariquidar as detected by LC-MS.

### P-gp inhibition restores inhibition of 26S proteasome and induction of apoptosis by Rapaprotin

Given that intracellular Rapaprotin-L levels are restored upon P-gp inhibition by tariquidar, we next assessed whether this leads to the same downstream effects in solid tumor cells as we have observed in MM cell lines, including inhibition of 26S proteasome activity and activation of the apoptotic pathway (Peng et al., 2025). We first determined the activity of the 26S proteasome whose inhibition is a hallmark of Rapaprotin activity. Western blot analysis of DLD-1 cell lysates revealed moderate accumulation of K48-linked polyubiquitinated proteins upon Rapaprotin treatment, which was markedly enhanced in the presence of tariquidar (Fig. 4A-B). This accumulation reflects impaired 26S proteasome function and confirms that restored intracellular Rapaprotin-L levels can recapitulate canonical 26S proteasome disruption.

**Figure 4.**
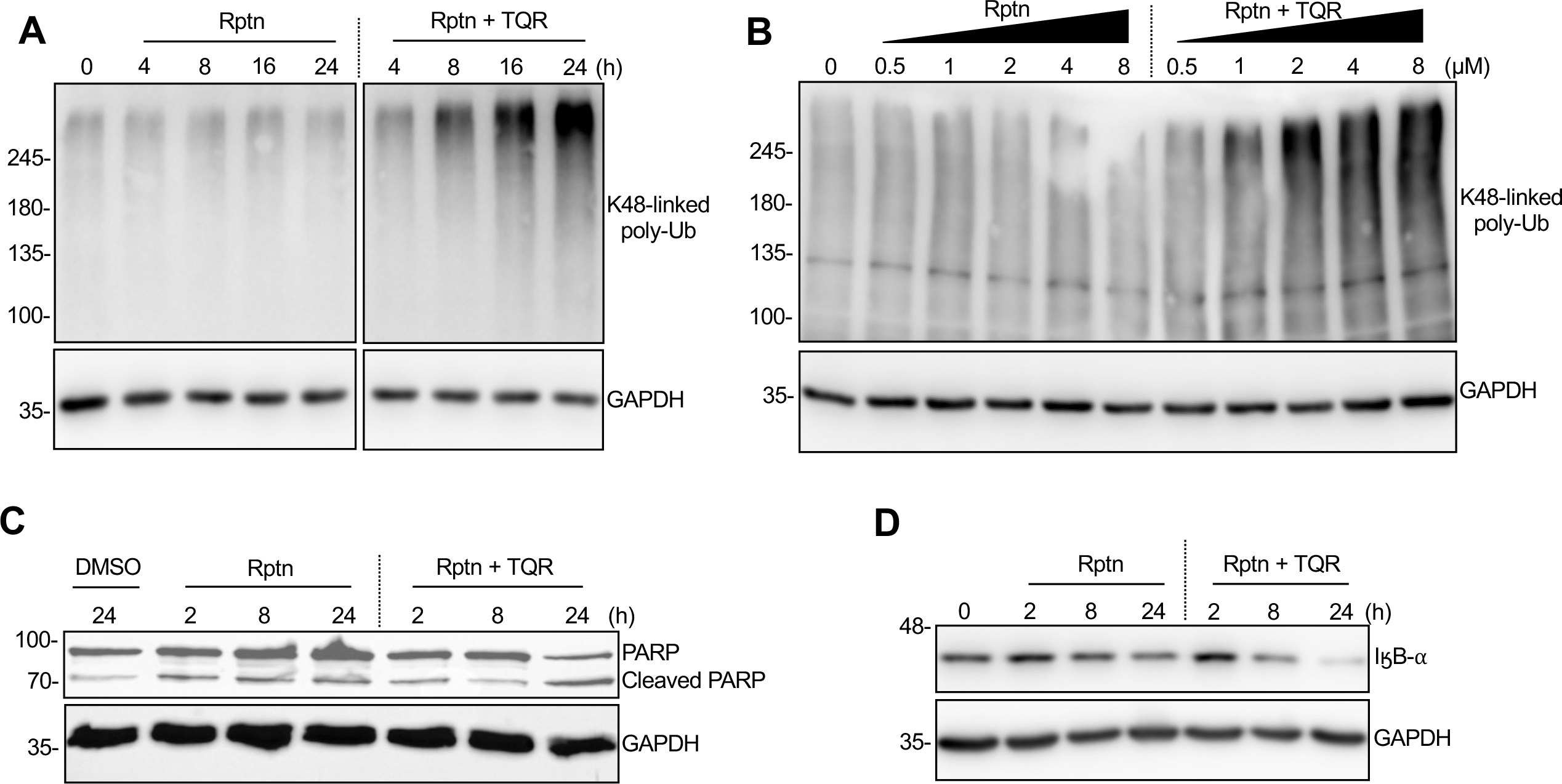
Rptn causes the accumulation of K48-linked Poly-Ub proteins and apoptosis in DLD-1 cell line when cotreated with TQR. (**A, B**) Anti-K48-linked Ub western blot showing both time-dependent (**A**) and dose-dependent (**B**) accumulation of polyubiquitinated proteins as a result of TQR and Rptn treatment in DLD-1 cells. (**C**) Western blot of DLD-1 cells treated with 0.2% DMSO, 500 nM Rapaprotin, and 500 nM Tariquidar. Data show modest PARP cleavage in the Rptn + TQR after 24 hours. (**D**) Western blot of IӄB-⍺ in DLD-1 cells with or without Rptn/TQR cotreatment. Data indicate diminished IӄB-⍺ levels after 24 hours upon concurrent treatment of Rptn + TQR.

To determine whether proteasome inhibition leads to cell death, we assessed apoptotic signaling by probing for cleavage of PARP by activated caspase. While Rapaprotin alone induced low levels of PARP cleavage in DLD-1 cells, co-treatment with tariquidar significantly enhanced PARP cleavage at 24 h, indicating synergistic induction of apoptosis (Fig. 4C). In addition, we examined the NF-κB stress response pathway by monitoring IκBα degradation. Co-treatment led to reduced IκBα levels relative to either agent alone, consistent with previous observations of proteotoxic stress triggering IκBα degradation and NF-κB pathway activation (Mathes et al., 2008; Tam et al., 2012) (Fig. 4D). Together, these results demonstrate that P-gp inhibition not only enhances Rapaprotin-L accumulation, but also reinstates its mechanistic capacity to disrupt proteostasis and induce cell death in solid tumor cells.

### Tariquidar sensitizes P-gp–high cancer cells to Rapaprotin

To determine whether the functional restoration of Rapaprotin activity translates into increased cytotoxicity, we performed viability assays using Rapaprotin in the presence and absence of tariquidar. We treated DLD-1, SW480, and HCT116 cells with varying concentrations of Rapaprotin and tariquidar and assessed viability after 72 h. Strong synergistic killing was observed in P-gp–high DLD-1 and SW480 cells, with Bliss synergy scores of 34.3 and 25.1, respectively (Fig. 5A-D). In contrast, Rapaprotin and tariquidar had an additive effect on HCT116 cell line which expresses a moderate level of P-gp (Bliss score: 9.9) (Fig. 5E-F). These results are consistent with the expression-dependent nature of P-gp–mediated resistance and further confirm that P-gp inhibition can reverse resistance in a cell-context–dependent manner. As control, we also determined if TQR is synergistic with other proteasome inhibitors such as Bortezomib in DLD-1 cells to confirm TQR synergism is unique to Rapaprotin. TQR exhibited no synergy with bortezomib (Fig. S1), suggesting that Rapaprotin is unique in its strong synergy with TQR in P-gp overexpressing cancer cells.

**Figure 5.**
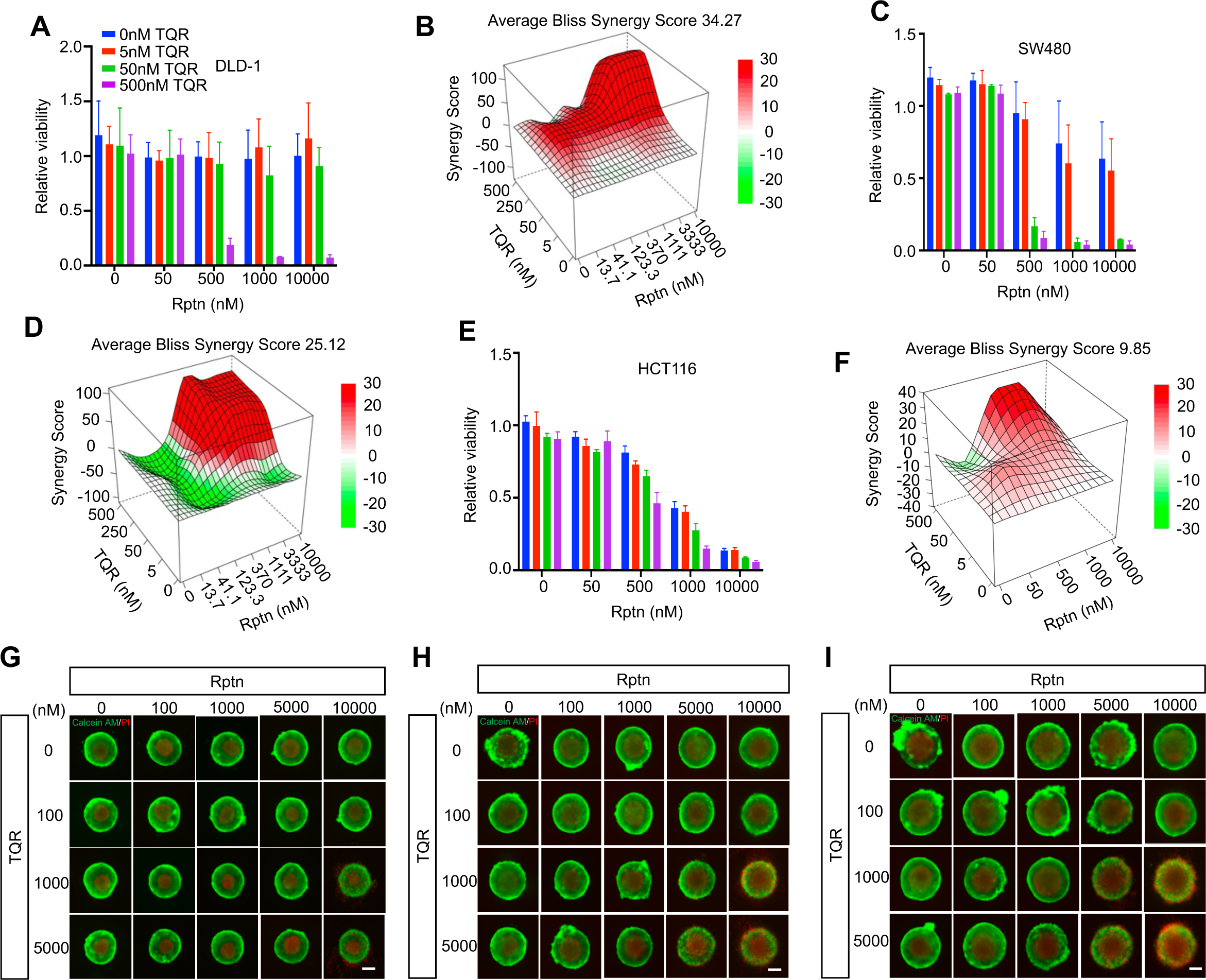
Rptn and TQR shows synergism in CRC cell lines that highly express P-gp. (**A-F**) Rptn was only sensitized by TQR in DLD-1 and SW480 cells. Alamar blue assay of DLD-1 (**A, B**), SW480 (**C, D**), and HCT116 (**E, F**) cell lines after 72 h treatment of Rptn with or without TQR was performed to determine the cell viability. Data were from three independent experiments and shown as mean ± SEM. (**G-I**) Cell staining of 3D-DLD-1 spheroids with designed concentration of Rptn and TQR. 3D-DLD-1 spheroids were treated for 3 days (**G**), 5 days (**H**), and 7 days (**I**), respectively, followed by staining with Calcium AM (green) and PI (red) to visualize the live and dead cells. Data were from three independent experiments.

To assess this synergy in a more physiologically relevant context, we utilized 3D tumor spheroid models. DLD-1 spheroids were treated with vehicle, Rapaprotin, tariquidar, or their combination. After 3, 5, and 7 days, spheroids were stained with propidium iodide (PI) to detect membrane-compromised, nonviable cells. Co-treatment with Rapaprotin and tariquidar led to marked PI uptake throughout the spheroid core over time, whereas monotherapies produced only peripheral staining or negligible cell death (Fig. 5G-I). This result indicates that tariquidar potentiates Rapaprotin’s activity not only in 2D culture but also in 3D tumor-like structures, raising the possibility of applying the Rapaprotin-tariquidar combination in the treatment of solid tumor cells that overexpress P-gp.

## Discussion

Using the PRISM drug screening platform, we identified P-glycoprotein (P-gp, ABCB1) as a key contributor to Rapaprotin resistance across a diverse set of human cancer cell lines. Resistance was most pronounced in solid tumor-derived lines, where high P-gp expression strongly correlated with reduced sensitivity to Rapaprotin. Specifically, DLD-1 and SW480 colorectal cancer cells, both characterized by high P-gp levels, were found to be resistant to Rapaprotin. Co-treatment with the P-gp inhibitor tariquidar restored Rapaprotin efficacy in these lines, directly implicating P-gp in mediating Rapaprotin efflux and resistance. In contrast, the NCI-H929 multiple myeloma cell line, originating from a hematologic malignancy, exhibited little to no P-gp expression and was highly sensitive to Rapaprotin. Other low P-gp–expressing lines, including A549, HeLa, HCT116, and MCF-7, showed variable levels of sensitivity, further reinforcing the inverse relationship between P-gp expression and Rapaprotin sensitivity. This differential expression of P-gp across tumor and tissue types may offer a therapeutic advantage. P-gp is basally expressed in tissues that form biological barriers such as the intestinal epithelium, blood– brain barrier, and renal tubules, where it functions to prevent intracellular accumulation of xenobiotics (Ahmed et al., 2022; Fu, 2013; Pilotto Heming et al., 2022). This physiological expression profile may protect healthy solid tissues from unintended Rapaprotin-L accumulation, likely reducing toxicity of Rapaprotin to normal tissues and organs. Meanwhile, liquid tumors, such as multiple myeloma, which typically express low levels of P-gp, remain accessible to Rapaprotin. In solid tumors where P-gp is upregulated, combination treatment with P-gp inhibitors such as tariquidar proves to be an effective strategy to overcome resistance, as shown by restored sensitivity to Rapaprotin in DLD-1 and SW480 cells. Interestingly, though a combination with TQR rendered HUVEC primary cells sensitive to Rapaprotin (Fig. S2), the IC_50_ value (4.6 µM) of Rapaprotin in the presence of 250 nM of TQR is over 10-fold higher than those of Rapaprotin in DLD-1 and SW480 cell lines in the presence of the same concentration of TQR (Fig. S2), offering a therapeutic window for the combination of Rapaprotin and TQR in the treatment of P-gp-high solid cancers without causing significant toxicity to normal cells.

P-gp is known to mediate the efflux of a variety of structurally diverse drugs and metabolites including rapamycin and FK506 (Saeki et al., 1993; Yacyshyn et al., 1996). Much to our surprise, Rapaprotin-L that bears a carboxylic acid group exhibited a high efflux ratio in MDCKII-MDR1 cells and accumulated intracellularly following P-gp inhibition with tariquidar, indicating that it is a high-affinity P-gp substrate. In contrast, the parent compound Rapaprotin with a scaffold similar to rapamycin and FK506 is not a substrate of P-gp and displayed a modest efflux ratio (∼4.3) by a P-gp–independent efflux mechanism. It is also interesting to note that while P-gp is a known promiscuous substrate transporter, the binding pocket is more accommodating to hydrophobic/positively charged molecules due to glutamate residues which reside deep in the substrate binding pocket (Aller et al., 2009; Martinez et al., 2014). Even more interesting is that it is the negatively charged Rapaprotin-L which is the substrate for P-gp and not its cyclic, uncharged parent compound Rapaprotin. There are a few examples of negatively charged P-gp substrates, the most prominent of which is Methotrexate in rats (Ogushi et al., 2017; Yokooji et al., 2007). We believe this efflux data firmly cements Rapaprotin-L as a rare instance of a negatively charged P-gp substrate. The superior efflux efficiency of negatively charged Rapaprotin-L by P-gp may be attributed to its five aromatic rings. The substrate-binding pocket of P-gp is enriched in aromatic amino acids such as phenylalanine and tyrosine, which facilitate strong π-π interactions with aromatic substrates (Aller et al., 2009).This abundance of aromatic groups in Rapaprotin-L likely compensates for the presence of a single negatively charged carboxyl group, allowing efficient recognition and export by P-gp. In contrast, the cyclic parent Rapaprotin is not a P-gp substrate, possibly because its aromatic moieties are conformationally constrained by the macrocyclic scaffold, limiting their ability to interact optimally with the aromatic residues within the P-gp binding pocket.

Inhibition of P-gp not only increased intracellular accumulation of Rapaprotin-L but also restored proteasome inhibition and pro-apoptotic activity of Rapaprotin. Co-treatment of tariquidar and Rapaprotin led to an accumulation of K48-linked polyubiquitinated proteins, consistent with disruption of the ubiquitin-proteasome system, along with increased PARP cleavage and IκBα degradation, indicative of apoptosis and cellular stress. In cell viability assays, strong synergy with Bliss synergy scores of 34.3 and 25.1 were observed between Rapaprotin and tariquidar in P-gp-overexpressing DLD-1 and SW480 cells, respectively. Importantly, in 3D spheroid models of DLD-1 cells, combination treatment increased membrane permeability, validating greater apoptotic activity in a physiologically relevant context. Such synergy between an inhibitor of P-gp and a P-gp substrate has seen success in several studies including clinical trials for doxorubicin, vinorelbine, and docetaxel in solid tumor models (Fox et al., 2015), docetaxel nano-formulations in cell culture (Kim et al., 2023), as well as vincristine in relapsed neuroblastoma models (Rosch et al., 2023). Together, these results suggest that the combination of Rapaprotin and tariquidar may allow for the expansion of potential clinical applications of Rapaprotin to a wider variety of cancer cells.

## Conflicts of Interest

The authors declare no competing financial interests.

## Acknowledgements

This work was supported in part by NIH (R01GM145793 and DP1CA174428), FAMRI, the Institute of Basic Biomedical Sciences of Johns Hopkins School of Medicine (JOL), and NCI (Cancer Center Support Grant: P30CA006973). The PRISM screening was supported by a sponsored research fund from Rapafusyn Pharmaceuticals. The NMR data was acquired using JEOL JNM-ECZL500R spectrometer at JHU Pharmacology NMR facility that was supported by NIH (S10OD034217). We thank Phillip R. Sanchez (NCATS) for advice on the 3D spheroid culture. We acknowledge Pharmaron (Ningbo, Zhejiang, China) for performing the P-gp substrate assay.

## Data Availability Statement

Data are available and will be provided upon request to the corresponding author.

## Supplementary information

**Supplementary Figure S1.**
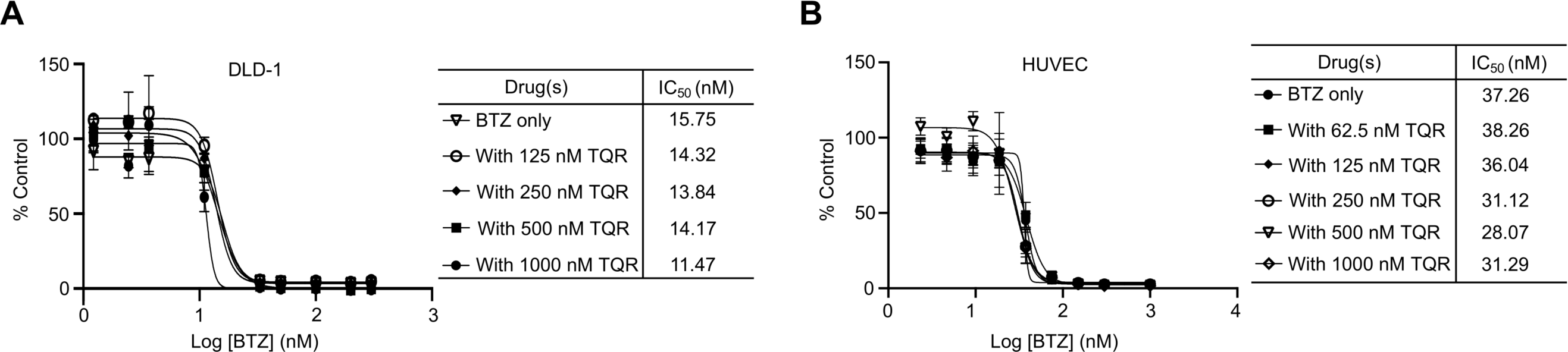
BTZ and TQR shows no synergism in DLD-1 (**A**) or HUVEC primary cells (**B**) in cell viability assays. Data were from three independent experiments and shown as mean ± SEM.

**Supplementary Figure S2.**
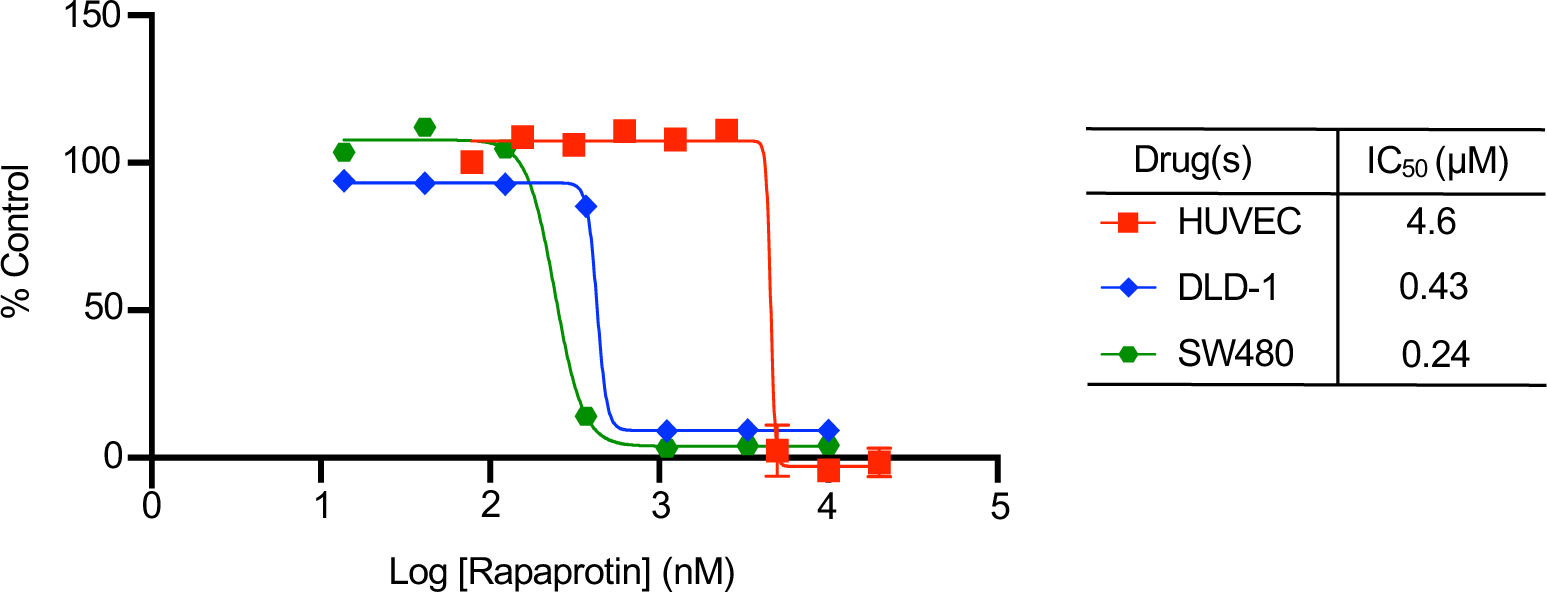
Combination of Rptn and TQR shows greater synergy in DLD-1 and SW480 cell lines than in primary HUVEC cells in cell viability assays. The concentration of TQR is fixed at 250 nM. Data were from three independent experiments and shown as mean ± SEM.

## Notes

### Competing Interest Statement

The authors have declared no competing interest.

